# Spaceflight Shifts in Community rRNA Copy Number in the Salivary Microbiome of Astronauts

**DOI:** 10.1101/2024.06.12.598653

**Authors:** Mark R. Williamson

## Abstract

The oral microbiome is stable, easily sampled, and can indicate disease. Using metagenomic data from GeneLab, I examined the effects of spaceflight on the human salivary microbiome using a composite community measure, average rRNA copy number. A higher copy number is associated with a faster growth rate and primary microbial succession. I found a significant increase in community weighted mean copy number between pre-spaceflight and during-spaceflight samples (p=0.0082). Furthermore, changes in abundance suggest a greater impact on individual species rather than phyla-level changes. Finally, a robustness analysis highlighted the importance of accurate copy number estimates and species-level identification.

**Graphical Abstract:** 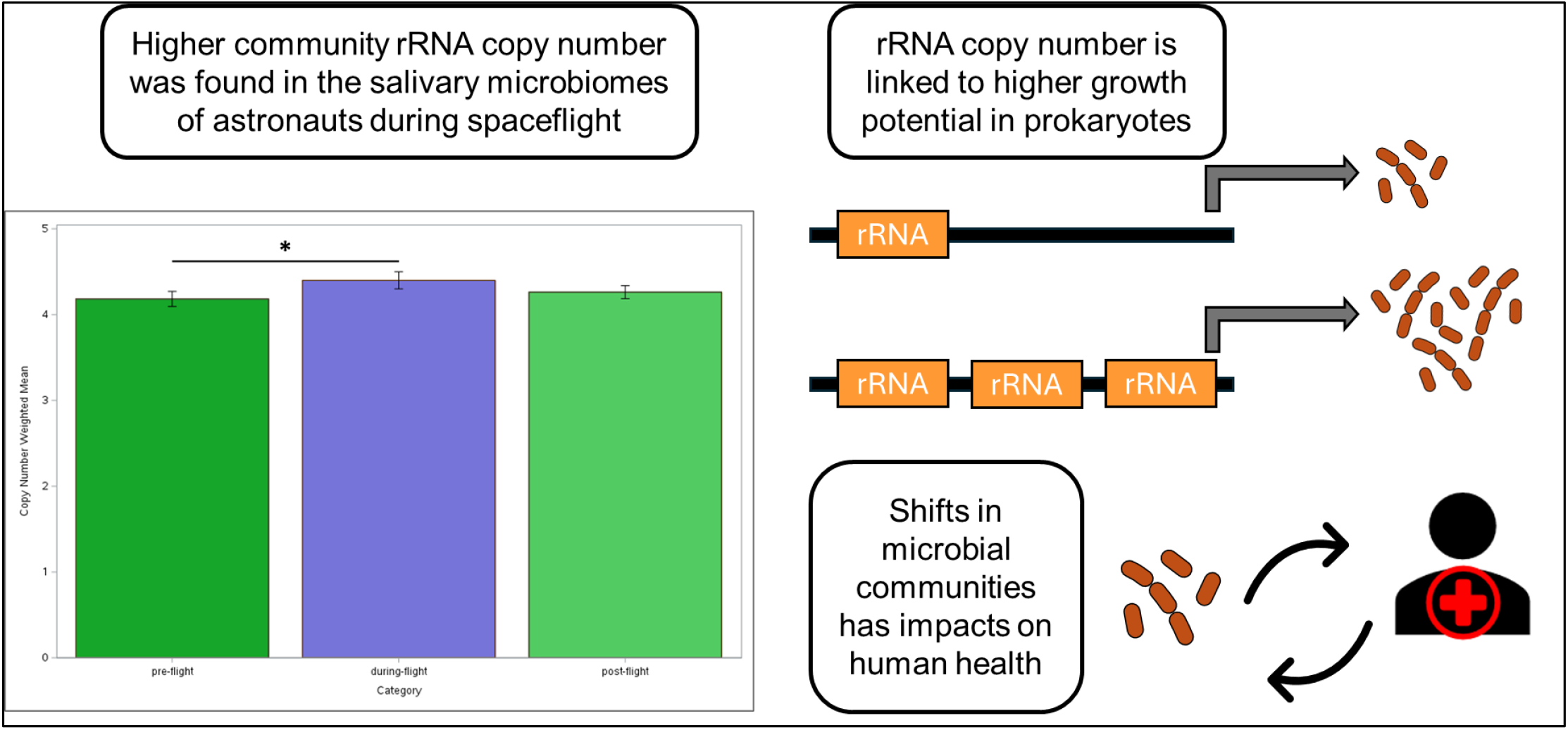

## Introduction

Ribosomal RNA copy number in prokaryotes is associated with life history traits [1]. Higher copy number is often found in copiotrophs and results in higher maximum growth rate, while lower copy number is more commonly found in oligotrophs. By studying community aggregated copy number, researchers found that average copy number decreases in microbial succession [2]. Similar results were found in a meta-analysis of bacterial primary succession [3] and specific cases such as the infant gut microbiome [4].

In the first stage of succession, where nutrients are more abundant and competition is limited, bacteria with higher maximum growth rates are expected to dominate. Higher rRNA copy number contributes by allowing for higher ribosome production, as ribosomal RNA makes up part of the physical structure of ribosomes, and thus directly links to protein synthesis. Indeed, experiments that manipulated the number of rRNA copies have found the maximum growth rate of the highest copies numbers under high nutrient conditions [5, 6]. However, observations of natural variation have not always replicated that [7]. In later stages of succession, the lower average copies number represents a shift to slower and more efficient growth. Because rRNA copies may be constitutively expressed in prokaryotes, rRNA transcription may be an ‘all-or-nothing’ event [8].

Less is known about primary succession in microbes compared to macroscopic communities [9], particularly in space. While some early work on how microbes interact in constructed space is available on NASA’s GeneLab portal [10-13], I was unable to find any research on primary succession. The components that make up the built environment in space are kept sterile during development [10]. However, microbes inevitably spread from populations that resisted the sterilization process or from the human microbiome.

The salivary microbiome is part of the larger oral microbiome and contains at least 500-700 species, though only a subset of around 300 are found within an individual [14-16]. Actinobacteria, Bacteroidetes, Firmicutes, Fusobacteria, and Proteobacteria represent the major phyla [14], although an increasing number of bacteria from the Candidate Phyla Radiation, or Patescibacteria, have been found [16]. The salivary microbiome is of interest as a predictor and cause of oral disease [15]. For example, changes in abundance in some species have been linked to periodontal disease [17]. It also has the potential to serve as a non-invasive biomarker for other diseases [14, 18]. The oral microbiome is established from birth, with much of the original taxa appearing to originate from maternal contact, though individualization begins early [19, 20].

An open question is how microbial community traits shift when the environment changes, especially in human microbiomes. As such, the effect of spaceflight and its associated stress on microbial community copy number is unknown. To answer that question, this study examines the changes in copy number of the human salivary microbiome pre-, during, and post-spaceflight. The hypothesis being tested is that the average community copy number will increase because the shift to space represents a perturbation of an otherwise stable environment. Therefore, bacterial species more primed to deal with shifts in the environments, like copiotrophs, will increase faster.

## Materials and Methods

### 1.1 Data Collection

The dataset OSD-280 (version 2) was obtained from GeneLab, NASA’s comprehensive space-related omics database. OSD-280 was originally collected as part of a study to examine salivary microbiome changes in astronauts across three time points: pre-spaceflight, during-spaceflight, and post-spaceflight [11]. Further details on how the data was collected in the study documents [21]. I used the processed 16S rRNA amplicon sequencing data, which included taxonomy and counts across samples, and included details on time points (pre-spaceflight, during-spaceflight, post-spaceflight) taken from the metadata file. From there, subset the count data to include only amplicon sequence variants (ASVs) with an average count of 10 or more across the 91 columns per sequence. This left 388 ASVs out of a total of 2286.

### 1.2 Species Identity and CN Estimation

The processed ASVs had taxonomic information that was classified down to genus, but not species. This is due to the difficulty of 16S amplicon sequencing in resolving species-level information. Therefore, to get species estimates, I obtained sequence information for the 388 ASVs provided in a FASTA file.

To compare sequences, I downloaded the local version of BLAST (blast+ version 2.14.1) and the 16S_ribsomal_RNA database. I used the operation blastn to run through the FASTA sequences, with a max target number of sequences set at 20. Using a custom Python script, I reduced the resulting output (hits) for each sequence. For an ASV, if there were one or more BLAST hits with a percent identity of 100% (perfect match), all perfect hits were kept. If there were no 100% percent ID hits, only the top identity was kept (ex. 98.5%).

From there, I manually went through the Ribosomal RNA Operon Copy Number Database (rrnDB) and pulled out the copy number for each hit from the previous step. If the exact species was not in the rrnDB, I took the average of the next lowest taxonomic ranking. This usually was resolved at genus. More rarely, I used a higher taxonomic ranking like family or order.

Then, using another custom Python script, I averaged copy number for each of the 388 ASVs. This meant sometimes averaging across multiple species that had hits for a single ASV. This resulted in every ASV having a copy number estimate.

### 1.3 Community Copy Number

The copy number estimate for each ASV was added to the dataset that included ASV counts across the 91 samples. Sequence abundance was calculated by dividing the number of counts for each sequence across the total number of sequences per sample. As in Nemergut et al, 2016 [2], I calculated the average copy number per column, also known as the community aggregated trait value of copy number. It is the product of the estimated operon copy number and the relative abundance for each OTU (in our case ASV), and then summing this value across all OTUs in that sample. This generated a community copy number estimate for each sample. I compared it to the approach in Guittar et al 2019 [4] as an additional confirmation of the correct approach. In that paper, community-level differences in a trait were measured using a community-weighted mean (CWM). A CWM is an average across OTUs of a microbial community, with a weighting correction that is based on each OTU’s relative abundance, where p is OTU abundance and x is the trait value, from i to the sample number (Equation 1). For consistency, I will use copy number community weighted mean (CN-CWM) for the rest of the report.

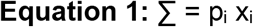

### 2.1 Data Analysis

I compared the CN-CWM across the three spaceflight categories (pre, during, post) by using a generalized linear model via PROC GLIMMIX in SAS. I modeled CN-CWM as a function of spaceflight category with a normal distribution (as Poisson and Negative binomial distribution were severely underdispersed). The resulting least squares means were compared with and without Tukey adjustment.

I ran several standard amplicon sequencing comparisons in R. Stacked bar plots were created with ggplot2 by combining ASV abundances by phylum and grouping across the three spaceflight categories. This was done for both raw abundances and copy-number corrected abundances. Principal components analysis was run using both raw and copy-number corrected abundances using the prcomp() function and plotted using the autoplot() and ggplotly() functions, color-coding samples by spaceflight category. Abundance and diversity metrics were calculated across time categories using the R packages ‘vegan’ and ‘picante’, for both raw and copy-number corrected abundances.

### 2.2 Robustness

To determine how much my approach to copy number estimation influenced the results, I sampled copy number from the theoretical range for each ASV and ran repeated linear models to test the robustness of the outcomes. First, using a custom Python script, I created a dictionary containing the taxonomic ranks generated from the GeneLab analysis for each ASV. From there, I created a dictionary for every entry in the rrnDB, which included copy number and taxonomic rank. Second, for each ASV in the first dictionary, I searched the rrnDB dictionary for the lowest taxonomic rank that matched the ASV taxonomy starting at genus and going up to domain. For the lowest taxonomic rank match, I appended a list of all copy number estimates for that rank to the ASV entry. The list of copy numbers represented—given no other information— the pool of known copy numbers the ASV could contain. Third, for those lists of potential copy numbers, I iterated through each ASV and pulled a random copy number from that list to serve as the copy number for that ASV. From there, I calculated the CWM for each sample. This was repeated for a total of 100 repetitions. Fourth, I took those CWM estimates, exported them to R, and using a loop, ran a generalized linear model (same parameters as the original model) for each repetition. The F-value and p-value were retained for each iteration and then tested against the initial F-value.

## Results

There was a significant difference in CN-CWM across spaceflight categories (F=4.71, p=0.0114). Mean copy number during spaceflight (4.42) was significantly higher than pre-spaceflight (4.22) and post-spaceflight (4.3). When using adjusted p-values, during spaceflight was still significantly higher than pre-flight, but not post-flight (Table 1). Given only three spaceflight categories, it may not be necessary to employ a *post-hoc* adjustment. However, even with a Tukey adjustment, mean copy number during spaceflight was still significantly higher than pre-spaceflight (Figure 1).

**Table 1:** Comparisons of spaceflight categories.

| Comparison | Estimate | Std. Error | t-value | p-value | adj. p-value |
| --- | --- | --- | --- | --- | --- |
| During vs. Post-spaceflight | 0.1258 | 0.06238 | 2.02 | **0.0468** | 0.1142 |
| During vs. Pre-spaceflight | 0.2041 | 0.06665 | 3.06 | **0.0029** | **0.0082** |
| Pre vs. Post-spaceflight | 0.07826 | 0.05795 | 1.35 | 0.1804 | 0.3714 |

**Figure 1:**
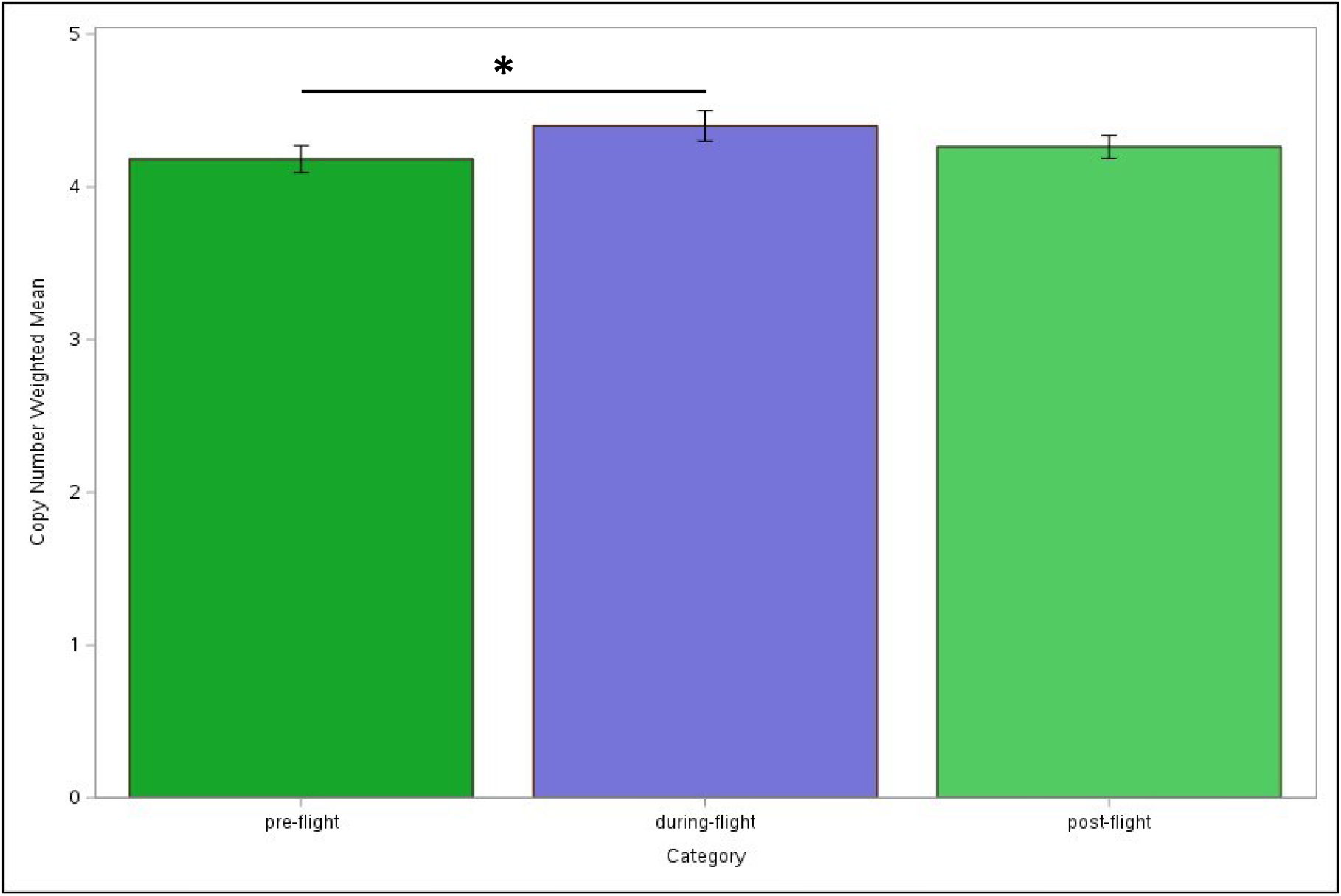
Mean copy number across spaceflight samples for the salivary microbiome of astronauts. The asterisk indicates a significant difference.

There were shifts in bacterial phyla abundance between spaceflight categories, especially the during-spaceflight samples. This was the case whether copy number correction was employed or not (Figure 2). The most notable shifts were an increase in Proteobacteria and decreases in Actinobacteria and Firmicutes during spaceflight, compared to pre- or post-spaceflight. Principal components analysis of microbiome analysis did not reveal any grouping across spaceflight categories. Samples were intermingled regardless of category, both for non-corrected and copy-number corrected abundances (Figure 3).

**Figure 2:**
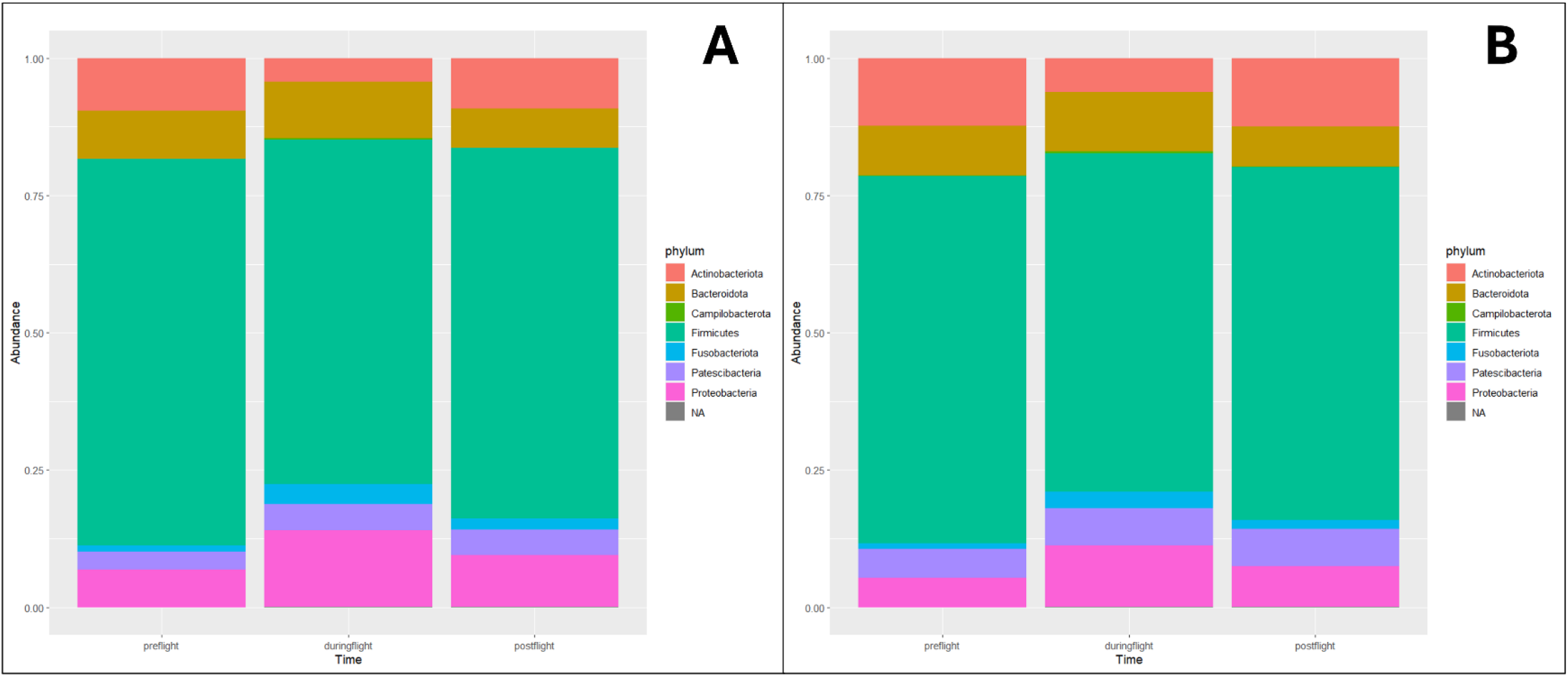
Bacterial phylum abundances across spaceflight samples. A) Raw abundances. B) Copy-number corrected abundances.

**Figure 3:**
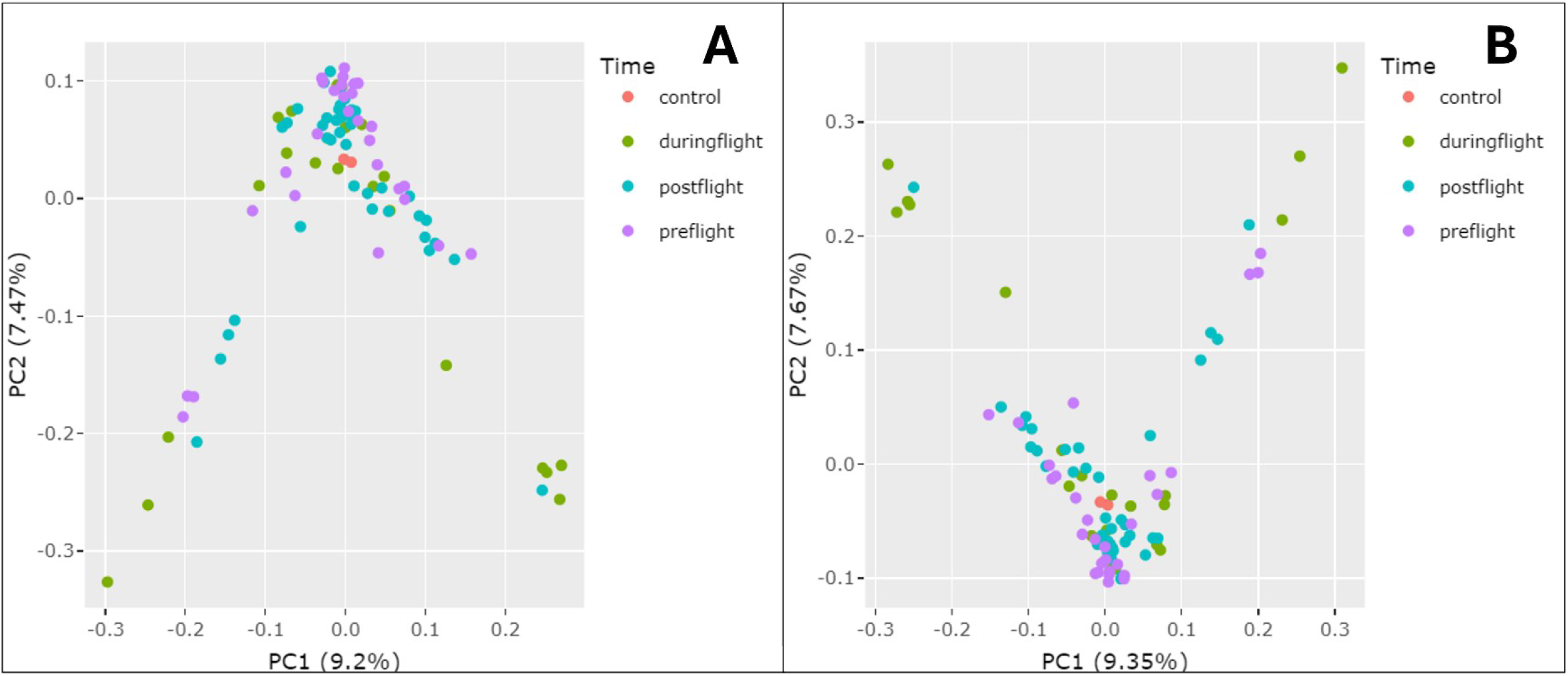
Principal components analysis across samples. A) Raw abundances. B) Copy-number corrected abundances.

The abundance and diversity of samples were fairly consistent across spaceflight categories. Pre-spaceflight samples had the highest mean abundance, as well as Simpson and Shannon diversity, while post-spaceflight had the lowest (Table 2). This suggests that spaceflight has a negative effect on diversity that may continue even after planetary reentry. For the test of robustness, a one-way Wilcox test using the 100 F-values generated from the sampling compared to the F-value from the initial analysis (constant value) was significant (p<0.0001). This indicates that the mean of the F-values from the random selection of iterations of CN was significantly different than the constant value. Most F-values were lower and gave non-significant regression tests (Figure 4).

**Table 2:** Abundance and diversity metrics.

| Time | Mean Abundance (Std) | Mean Simpson (Std) | Mean Shannon (Std) |
| --- | --- | --- | --- |
| Control | 23.5 (2.12) | 0.46 (0.12) | 1.24 (0.36) |
| Pre-spaceflight | 158.6 (35.3) | 0.88 (0.08) | 3.18 (0.64) |
| During spaceflight | 150.8 (32.5) | 0.87 (0.08) | 2.98 (0.46) |
| Post spaceflight | 146.3 (48.7) | 0.85 (0.10) | 2.85 (0.58) |

**Figure 4:**
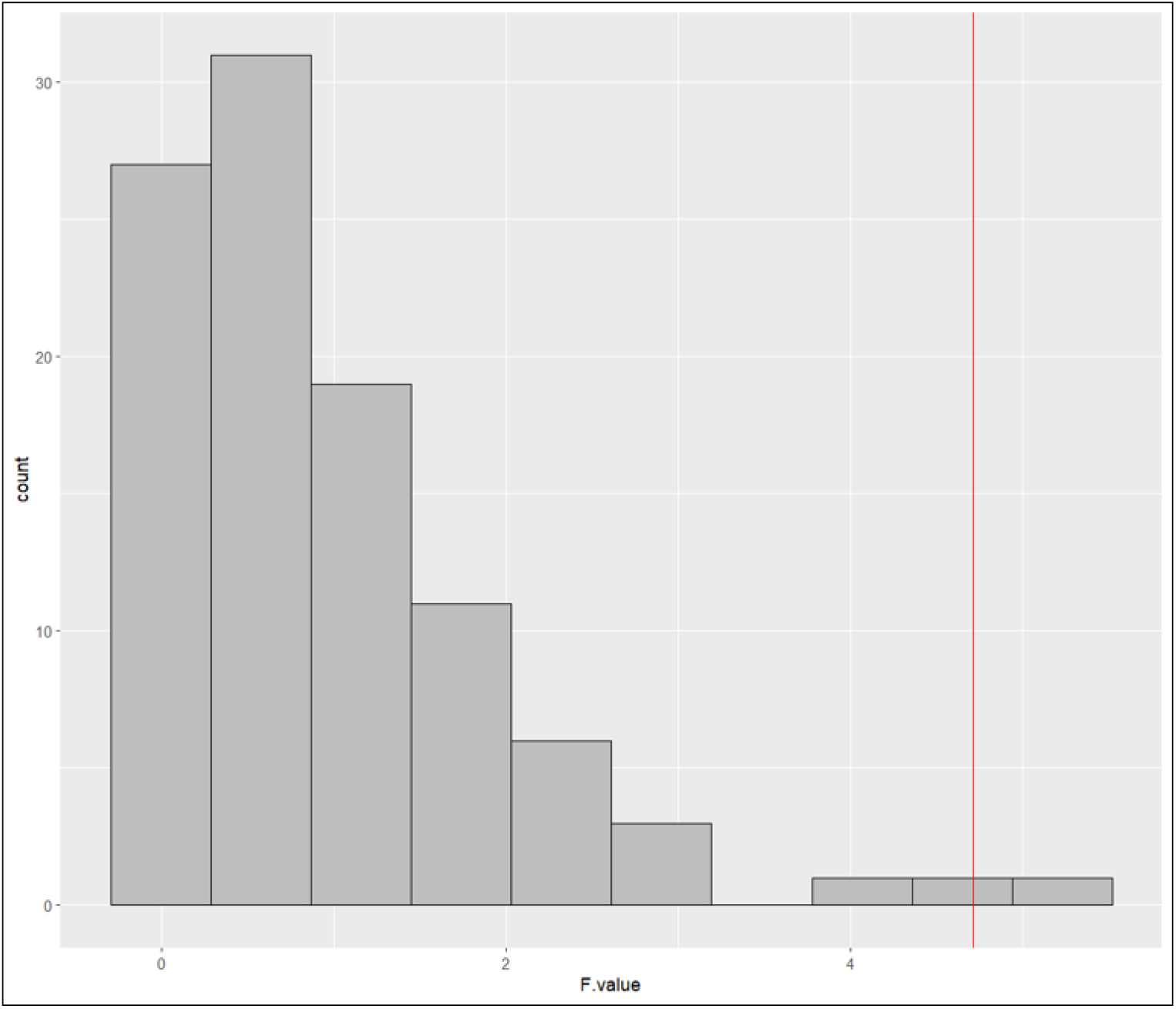
Distribution of F-values from robustness analysis. The red line is the F-value of the main analysis.

## Discussion

In comparing mean copy number of salivary microbiome samples across categories (pre-, during-, and post-flight), I found a significant increase in the during-flight category. The shift was driven by increases in Proteobacteria and decreases in Actinobacteria and Firmicutes. Species assemblages were not clustered across categories and abundance and diversity decreased during spaceflight. In a robustness study that randomly selected CN, there was a significant difference in the observed and simulated F-values, highlighting the sensitivity of current CN estimations.

The results from Firmicutes were somewhat surprising. It was reasonable for Proteobacteria to increase, as mean copy number for the phylum is 5.5 (median=6). It also made sense for Actinobacteria to decrease, with an average copy number of 3.2 (median=3.0). However, Firmicutes has one of the highest average copy numbers of any bacteria phylum, at 6.9 (median=6). Its decrease during flight suggests more of an emphasis on individual species in contributing to CN-CWM than the phylum as a whole.

To examine this further, I averaged the abundance of individual ASVs across the three spaceflight categories and checked the changes in abundance between pre- and during-spaceflight. The top 10 ASVs for increased abundance had an average CN of 4.58 and were dominated by Proteobacteria, three being from the genus Haemophilus. The top 10 ASVs for decreased abundance had an average CN of 3.82 and consisted mostly of Firmicutes, five being from the genus Streptococcus.

The fact that a significant shift in CN-CWM was detected during spaceflight for the salivary microbiome is interesting. The oral microbiome in general is resistant to local changes such as diet [16]. The microbiome also resists external changes to the human host, like short-term hospitalization [18]. However, oral hygiene and antibiotic use have major impacts. Importantly, the oral microbiome appears to resist external colonization under normal circumstances [16], so the lower microbe density found in space may affect colonization pressure.

However, the direction of the shift, increased CN-CWM, is in line with general studies on growth rate, which generally have found an increase in bacterial growth in space [22]. This would be in line with increased copy number because faster-growing species tend to have higher copy number. Bacterial studies in space and simulated microgravity have shown higher population growth compared to ground controls in many studies, mostly in liquid cultures [23]. However, in other cases, the growth was slower in microgravity [24]. Broadly speaking, the effects of microgravity on metabolism depend on the specific organism [25]. This may be due to non-mobile species staying in suspension rather than settling to the bottom, thus showing increased growth, while mobile species do not show increased growth because there is no major change in distribution [26].

Bacterial responses in the salivary microbiome and other human microbiomes can have environmental effects. Microbes found in space environments like the ISS come from species that either survive the disinfection protocols for spacecraft or from the astronauts themselves [27]. The major stresses on microbial life in space come from microgravity and radiation [28]. Microbes change in response to the stresses of the space environment, including metrics that can harm human health such as biofilm formation, virulence, and antibiotic resistance [25, 28, 29]. Biofilm formation increases bacterial resistance to stresses and can contribute to chronic diseases in humans, such as in the oral cavity [28]. Therefore, it is important to continue exploring the dynamics of microbial communities in the space environment.

A major shortcoming of this study was that accurate copy number estimates were not known for all species, especially since the lack of resolution in 16S amplicon sequencing precluded accurate species identity, let alone isolate count. While we did find a significant result using the best average of CN that was available, the robustness study showed that a wide landscape of potential CN values and that random selection resulted in few simulations where there was a significant difference between spaceflight categories. More accurate studies, with whole genome sequencing or similar expansions, and the growth of CN estimates for more species and isolates are needed to fill in this shortcoming.

Future work should focus on more spaceflight microbiome data. Ultimately, it would be ideal to perform succession studies on microbes in spaceflight environments as a broader test of the hypothesis that spaceflight succession results in an increase in community mean copy number. In conclusion, bacterial copy number is related to growth rate and has been shown in this analysis to increase during spaceflight. As humans continue to live and work in space, microbes will continue to be a part of that endeavor. Understanding more about microbial growth mechanisms will help humans keep their bodies and spaceflight environment healthy by addressing the shifts in the human microbiome and the spread of health-impacting external microbe communities.

## Acknowledgements

The author would like to thank Dr. Brian Darby for his helpful feedback and suggestions.

## Data Availability Statement

- Initial datasets from OSD-280, Salivary microbiome sequencing of astronauts, can be found at https://osdr.nasa.gov/bio/repo/data/studies/OSD-280.
- The following datasets can be found at the GitHub repository: <u>SalivaryMicrobiomeCN</u>
  - **Amplicon_ASVs_reduced_CN.csv:** copy number estimates, based on the Ribosomal RNA Copy Number Database, for each ASV used in the study
  - **Salivary Microbiome Abundance CN corrected Table.csv:** the copy number corrected abundance tables across samples for each ASV used in the study
  - **Salivary_Microbiome_CN_final_ds.csv:** the final dataset, with CWM for each sample

## References

1. Klappenbach, J.A., J.M. Dunbar, and T.M. Schmidt, rRNA operon copy number reflects ecological strategies of bacteria. Appl Environ Microbiol, 2000. 66(4): p. 1328–33.

2. Nemergut, D.R., J.E. Knelman, S. Ferrenberg, T. Bilinski, B. Melbourne, L. Jiang, … A.R. Townsend, Decreases in average bacterial community rRNA operon copy number during succession. ISME J, 2016. 10(5): p. 1147–56.

3. Ortiz-Alvarez, R., N. Fierer, A. de Los Rios, E.O. Casamayor, and A. Barberan, Consistent changes in the taxonomic structure and functional attributes of bacterial communities during primary succession. ISME J, 2018. 12(7): p. 1658–1667.

4. Guittar, J., A. Shade, and E. Litchman, Trait-based community assembly and succession of the infant gut microbiome. Nat Commun, 2019. 10(1): p. 512.

5. Condon, C., D. Liveris, C. Squires, I. Schwartz, and C.L. Squires, rRNA operon multiplicity in Escherichia coli and the physiological implications of rrn inactivation. J Bacteriol, 1995. 177(14): p. 4152–6.

6. Gyorfy, Z., G. Draskovits, V. Vernyik, F.F. Blattner, T. Gaal, and G. Posfai, Engineered ribosomal RNA operon copy-number variants of E. coli reveal the evolutionary trade-offs shaping rRNA operon number. Nucleic Acids Res, 2015. 43(3): p. 1783–94.

7. Valdivia-Anistro, J.A., L.E. Eguiarte-Fruns, G. Delgado-Sapién, P. Márquez-Zacarías, J. Gasca-Pineda, J. Learned, … V. Souza, Variability of rRNA Operon Copy Number and Growth Rate Dynamics of Bacillus Isolated from an Extremely Oligotrophic Aquatic Ecosystem. 2016. 6.

8. Williamson, M.R., Evolutionary Ecology of Ribosomal RNA Copy Number: Studies in Soil Prokaryotes and Nematode Communities, in Biology. 2019, University of North Dakota: Grand Forks.

9. Fierer, N., D. Nemergut, R. Knight, and J.M. Craine, Changes through time: integrating microorganisms into the study of succession. Res Microbiol, 2010. 161(8): p. 635–42.

10. La Duc, M.T., P. Vaishampayan, H.R. Nilsson, T. Torok, and K. Venkateswaran, Pyrosequencing-derived bacterial, archaeal, and fungal diversity of spacecraft hardware destined for Mars. Appl Environ Microbiol, 2012. 78(16): p. 5912–22.

11. Urbaniak, C., H. Lorenzi, J. Thissen, C. Jaing, B. Crucian, C. Sams, … S. Mehta, The influence of spaceflight on the astronaut salivary microbiome and the search for a microbiome biomarker for viral reactivation. Microbiome, 2020. 8(1): p. 56.

12. Venkateswaran, K., P. Vaishampayan, J. Cisneros, D.L. Pierson, S.O. Rogers, and J. Perry, International Space Station environmental microbiome - microbial inventories of ISS filter debris. Appl Microbiol Biotechnol, 2014. 98(14): p. 6453–66.

13. Ritchie, L.E., S.S. Taddeo, B.R. Weeks, F. Lima, S.A. Bloomfield, M.A. Azcarate-Peril, … N.D. Turner, Space Environmental Factor Impacts upon Murine Colon Microbiota and Mucosal Homeostasis. PLoS One, 2015. 10(6): p. e0125792.

14. Acharya, A., Y. Chan, S. Kheur, L.J. Jin, R.M. Watt, and N. Mattheos, Salivary microbiome in nonoral disease: A summary of evidence and commentary. Arch Oral Biol, 2017. 83: p. 169–173.

15. Nasidze, I., J. Li, D. Quinque, K. Tang, and M. Stoneking, Global diversity in the human salivary microbiome. Genome Res, 2009. 19(4): p. 636–43.

16. Wade, W.G., Resilience of the oral microbiome. Periodontol 2000, 2021. 86(1): p. 113–122.

17. Diao, J., C. Yuan, P. Tong, Z. Ma, X. Sun, and S. Zheng, Potential Roles of the Free Salivary Microbiome Dysbiosis in Periodontal Diseases. Front Cell Infect Microbiol, 2021. 11: p. 711282.

18. Cabral, D.J., J.I. Wurster, M.E. Flokas, M. Alevizakos, M. Zabat, B.J. Korry, … P. Belenky, The salivary microbiome is consistent between subjects and resistant to impacts of short-term hospitalization. Sci Rep, 2017. 7(1): p. 11040.

19. Mason, M.R., S. Chambers, S.M. Dabdoub, S. Thikkurissy, and P.S. Kumar, Characterizing oral microbial communities across dentition states and colonization niches. Microbiome, 2018. 6(1): p. 67.

20. Lif Holgerson, P., A. Esberg, A. Sjodin, C.E. West, and I. Johansson, A longitudinal study of the development of the saliva microbiome in infants 2 days to 5 years compared to the microbiome in adolescents. Sci Rep, 2020. 10(1): p. 9629.

21. Venkateswaran, K., C. Urbaniak, H. Lorenzi, J. Thissen, C. Jaing, B. Crucian, … S. Mehta, Salivary microbiome sequencing of astronauts. 2020, NASA Open Science Data Repository.

22. Horneck, G., D.M. Klaus, and R.L. Mancinelli, Space microbiology. Microbiol Mol Biol Rev, 2010. 74(1): p. 121–56.

23. Benoit, M.R. and D.M. Klaus, Microgravity, bacteria, and the influence of motility. Advances in Space Research, 2007. 39(7): p. 1225–1232.

24. Huang, B., D.G. Li, Y. Huang, and C.T. Liu, Effects of spaceflight and simulated microgravity on microbial growth and secondary metabolism. Mil Med Res, 2018. 5(1): p. 18.

25. Sharma, G. and P.D. Curtis, The Impacts of Microgravity on Bacterial Metabolism. Life (Basel), 2022. 12(6).

26. Baker, P.W. and L. Leff, The Effect of Simulated Microgravity on Bacteria from the Mir Space Station. Microgravity-Science and Technology, 2004. XV.

27. Moissl-Eichinger, C., C. Cockell, and P. Rettberg, Venturing into new realms? Microorganisms in space. FEMS Microbiol Rev, 2016. 40(5): p. 722–37.

28. Rosenzweig, J.A., O. Abogunde, K. Thomas, A. Lawal, Y.U. Nguyen, A. Sodipe, and O. Jejelowo, Spaceflight and modeled microgravity effects on microbial growth and virulence. Appl Microbiol Biotechnol, 2010. 85(4): p. 885–91.

29. Salavatifar, M., S.M. Ahmadi, S.D. Todorov, K. Khosravi-Darani, and A. Tripathy, Impact of Microgravity on Virulence, Antibiotic Resistance and Gene Expression in Beneficial and Pathogenic Microorganisms. Mini Rev Med Chem, 2023. 23(16): p. 1608–1622.

